# Heritable environments: bias due to conditioning on a collider in models with polygenic scores

**DOI:** 10.1101/2020.11.02.364539

**Authors:** Evelina T. Akimova, Richard Breen, David M. Brazel, Melinda C. Mills

## Abstract

The application of polygenic scores has transformed our ability to investigate whether and how genetic and environmental factors jointly contribute to the variation of complex traits. Modelling the complex interplay between genes and environment, however, raises serious methodological challenges. Here we illustrate the largely unrecognised impact of gene-environment dependencies on the identification of the effects of genes and their variation across environments. We show that controlling for heritable covariates in regression models that include polygenic scores as independent variables introduces endogenous selection bias when one or more of these covariates depends on unmeasured factors that also affect the outcome. This results in the problem of conditioning on a collider, which in turn leads to spurious associations and effect sizes. Using graphical and simulation methods we demonstrate that the degree of bias depends on the strength of the gene-covariate correlation and of hidden heterogeneity linking covariates with outcomes, regardless of whether the main analytic focus is mediation, confounding, or gene × covariate (commonly gene × environment) interactions. We offer potential solutions, highlighting the importance of causal inference. We also urge further caution when fitting and interpreting models with polygenic scores and non-exogenous environments or phenotypes and demonstrate how spurious associations are likely to arise, advancing our understanding of such results.

## Introduction

The importance of understanding the joint contributions of genetic and environmental variation underlying complex traits is widely recognised. The rise of polygenic scores has resulted in a surge of studies investigating the mediating and moderating roles of environments along with genetic confounding^1^ (i.e., whether genes confound associations between environments or phenotypes). Yet, disentangling the relative importance of polygenic scores and environmental covariates is difficult; various methodological concerns have been raised, including but not limited to the power and predictive accuracy of polygenic scores^2–4^ and the non-exogenous nature of environmental exposures and their consequences.^5–7^ Moreover, genes and environments do not operate independently, necessitating greater scrutiny of both conventional models and new methods addressing gene-environment-trait correlations.^8; 9^ Here we address further methodological problems arising from gene-environment correlations that have gone largely unrecognised yet make identification of causal effects and the accurate estimation of associations even more challenging.

We illustrate how the presence of genetic predispositions associated with exposures to an environment (or phenotype in cases of genetic confounding) introduces *endogenous selection bias* in a regression analysis. Heritable covariates in regression models with polygenic scores are endogenous variables, and this can give rise to the problem of conditioning on a collider. Collider bias is an important statistical problem that destabilises regression models and it can arise for a variety of reasons, including sample selection and attrition.^10^ We demonstrate that collider bias is likely to occur not only in genetic association studies but also in other analyses where polygenic scores are included, regardless of whether the main focus is mediation, confounding, or gene × environment (G×E) interaction.

Moreover, the issue we describe here is linked to a growing body of literature showing the heritability of environments known as gene-environment correlation (rGE). To date, discussion of the methodological implications of these findings has focussed on the implications for G×E interaction studies (e.g. Dudbridge & Fletcher^7^). However, if both genetic and environmental covariates are included in a statistical model, gene-environment correlations may lead to spurious estimates and effect sizes. Understanding the mechanisms that generate these dependences is crucial for how we interpret such results. We emphasise the conceptual differences between passive, evocative, and active gene-environment correlations^11^ and potential sources of endogeneity of environmental covariates.

We use a graphical approach to demonstrate these methodological problems, illustrated by simulations. We then discuss the consequences of the bias in linear models and offer some potential solutions. The problems outlined here are relevant for making both causal and non-causal claims, with serious implications for the interpretation of results.

### Endogenous selection bias

The notion of *endogenous selection bias* arises from the broader concept of *selection bias*. While the term *selection bias* is very widely used, ^12^ *endogenous selection bias* commonly arises in analyses in which we adjust for an endogenous variable – that is, a variable caused by other, unmeasured variables which also affect the outcome. In this case, bias arises through the adjusting variable operating as a collider. Collider bias was demonstrated in Day et al.^13^ in the context of genetic association studies where such bias led to false-positive and biologically implausible associations between GWAS significant SNPs for height and sex once the respondent’s height was adjusted for. The bias arose because the respondent’s height is a collider variable since it is a direct product of another covariate (SNPs of height) and an outcome (sex).

In what follows we consider a situation in which genes and environment are correlated (for reasons discussed below) and the environmental variable(s) is affected by variables not measured in the study and which also affect the outcome. We discuss the consequences of the resulting collider bias for both additive and G×E interaction models.

### Additive models with polygenic scores

The first type of model we consider is the rather straightforward design where polygenic scores and environmental covariates (or phenotypes if they are used as covariates) are jointly included as a set of predictors for an outcome of interest. Such models are intended to reveal whether genetic influences confound associations between environments and outcomes or whether environments are mediators of the link between genetic variants and phenotypes. Examples include linking health disparities with socio-economic outcomes such as the relationship between attention-deficit hyperactivity disorder (ADHD), its polygenic prediction, and IQ on educational outcomes among teenagers (e.g. Stergiakouli et al.^14^). Another example is studies of labour market outcomes predicted by educational measures (e.g., grades, years of education) along with an educational attainment polygenic score (e.g. Ayorech et al.^15^; Papageorge & Thom^16^). The variation of exam scores in relation to school types and polygenic prediction of education is another instance (e.g. Smith-Woolley et al.^17^).

All of these studies are similar regarding the nature of environmental variables – they are not exogenous, being the direct or indirect products of polygenic scores which are also included into the models. Dependence of these covariates could arise through the inclusion of a phenotypic variable – the scenario that is prevalent in genetic confounding studies. For example, the polygenic score for ADHD is directly linked to the diagnosis of ADHD and is a phenotype in the associated GWAS studies. Dependences could also arise due to active and evocative selection in environments. Applying the polygenic prediction of educational attainment as an example, we see that it contributes to the variation of school grades^17^ which likely reflects active rGE (i.e., children selecting their environments for genetically influenced reasons). It could also be linked to school type since parents adjust their educational choices for children depending on their child’s characteristics which are partially due to genetic differences, reflecting evocative rGE (i.e., the child indirectly shapes the environment via the reaction of parent’s to the child’s behaviour).^18; 19^ Therefore, non-exogenous environments of this type vary depending on the values of polygenic scores. Such dependencies accompanied by the presence of hidden heterogeneity linking environments with outcomes will result in endogenous selection bias, which we describe now.

Whether the aim is to address genetic confounding or to reveal mediation, models of this kind include polygenic scores and environments as predictors of an outcome of interest. The simple case is illustrated in Figure 1A. The polygenic score, G, has an independent association with the outcome, Y, along with an indirect path through the environment, E. The exclusion from the model of the environmental covariate, E, results in the estimation of the total effect of G on Y, while the exclusion of G and the inclusion of E produces the association between E and Y, confounded by unobserved G.

**Figure 1.**
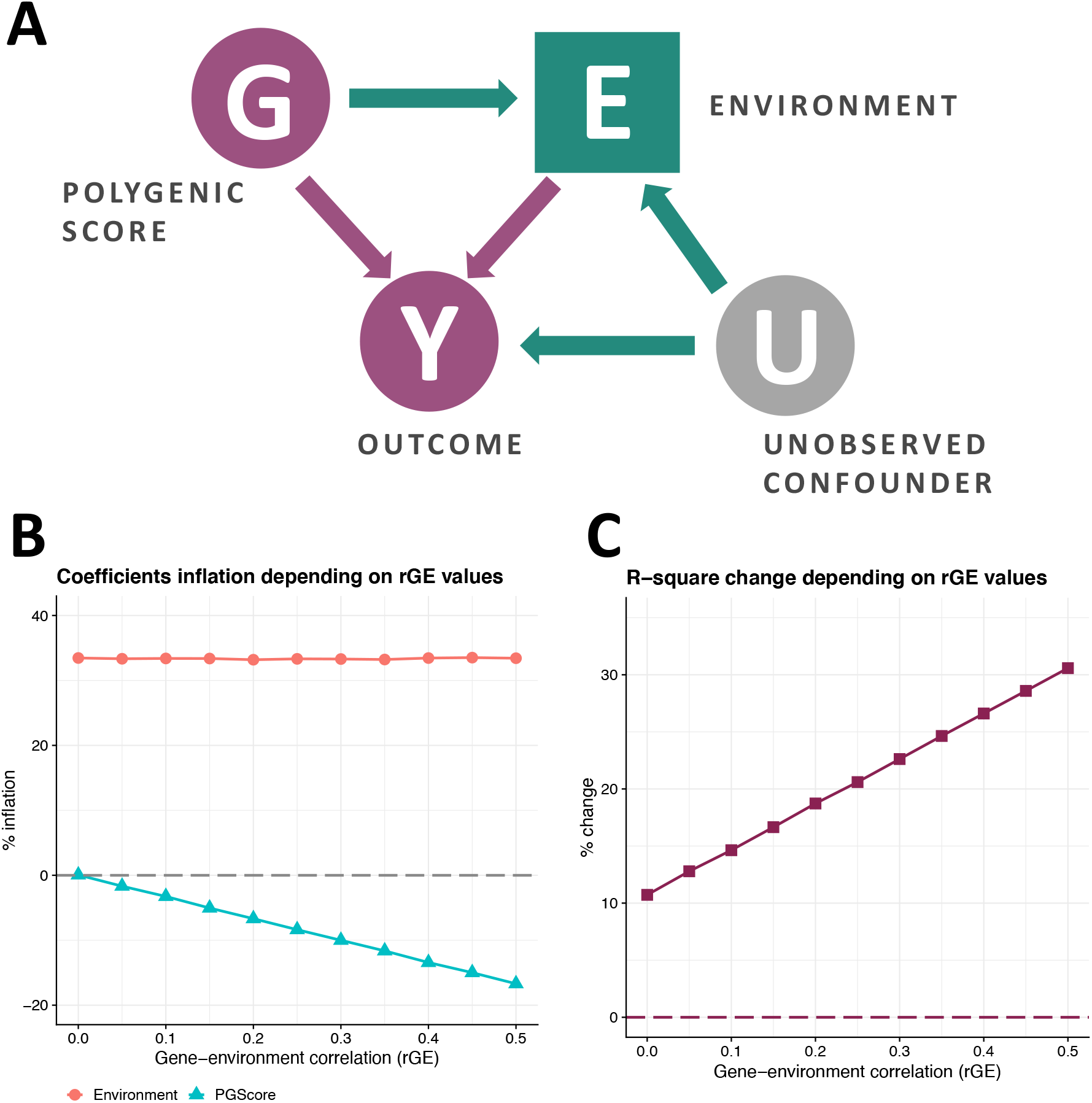
Collider bias in polygenic gene-environment models. Panel A. Schematic diagram of the collider bias which occurs between polygenic score, environment, and outcome in cases of gene-environment interdependence. Dark purple circles represent variables, unobserved confounders are shown in grey circles, collider variables are indicated by squares. By adding E into the model with the polygenic score G, we make E a collider. A collider that is not conditioned on, blocks the path between its sources (G and U); once a collider is controlled for, the path is opened as indicated by green nodes. Panel B. Spurious regression estimates for polygenic score and environment from the series of OLS simulations reflecting the range of gene-environment interdependence. Collider bias due to positive values of gene-environment correlation and the presence of uncontrolled confounder, which is positively correlated with covariate and outcome, results in deflation of polygenic score estimates. Estimates of the environmental effect are upwardly biased but are not affected by the gene-environment correlation. Panel C. R-squared inflation plot from the series of OLS simulations; collider bias results in inflated values of explained variance statistics. R-squared statistics for the model with endogenous covariate and polygenic score includes not only the true share of the variance in Y explained by G and E (baseline estimate indicated by 0), but also the elements of variance that are due to gene-environment correlation and confounder(s), U.

The challenges for the model in Figure 1A are to produce reliable estimates of the direct effects from G to Y and from E to Y in the face of confounding by the unmeasured U. Since the focus of our paper is not related to the issues of polygenic prediction *per se*, we do not include a discussion on the sources of bias between G and Y caused by confounders that are likely to arise due to assortative mating or population stratification, which have been amply explored elsewhere.^20^ Here, we focus on the role of confounders of the link between environment and outcome.

The most important problem arising from the presence of unmeasured factors causing E and Y is that of non-exogenous environments. Factors such as (but not limited to) socio-economic conditions, parental characteristics, health policies, cultural norms, and neighbourhood characteristics could cause E and Y to be correlated and, unless included in the model, they will be jointly present in the error structure of both variables. The issue is further problematic because the confounding can be driven by both observed and unobserved factors. Hence, even an extensive set of controls would not necessarily yield unbiased estimates if substantial confounding on unobservables remained unaddressed.

Moreover, unobserved confounder(s), U, linking E and Y biases not only their association, but also the estimate of the direct effect of G on Y. This is driven by the fact that E is now a collider since it is a product of both G and U, as indicated by the arrows from U to E and G in Figure 1A. It is known that if we do not control for a collider variable the path between its sources is blocked; however, once a collider is included in the set of covariates, the associated path is now open.^21^ Accordingly, conditioning on E introduces a new path from G to E to Y through U: this is the green path denoted in Figure 1A. This path is the source of the collider bias in these models.

To further illustrate this bias, we conducted a series of simulations of the simple linear model from Figure 1A. We considered the presence of direct effects from G to Y and E to Y, allowed the G-E correlation to vary from 0 indicating no heritability to 0.5 indicating a highly heritable covariate E, and included an uncontrolled confounder, U, which is positively correlated with both E and Y at a fixed value. Figure 1B illustrates the deviations of coefficients from the true simulated values. Notably, the presence of G-E correlation and an omitted confounder, U, where both U-E and U-Y associations are positive, results in the deflation of polygenic score estimates and inflation of environmental coefficients. Deflation of the G-Y association is greater with higher values of the G-E correlation, while the models without this association produced results free of collider bias. The path coefficient from E to Y is biased regardless of the strength of the G-E correlation reflecting the role of the omitted confounder, U, as a source of confounding of this path.

It is also possible to exemplify the source of inflation of the G-Y association by considering examples from the existing literature. For instance, Papageorge and Thom^16^ regress the educational attainment polygenic score and years of schooling on standardised job tasks. If we take models on nonroutine analytic and interactive tasks (where the association between polygenic score and outcome is positive), we see that the inclusion of educational controls results in about 70% smaller polygenic score coefficients. It is likely that such a change is largely attributable to the extended set of educational controls, which includes both parental and respondent educational attainment. However, if we consider a moderate strength of association between the genetic score and respondent’s years of schooling along with additional assumptions about unobserved confounders linking educational attainment and the type of job tasks, we can show that around 15-20% of the polygenic score coefficient decrease is plausibly due to collider bias, following the derivations we include in the Supplementary Data.

We also show in Figure 1C that the described bias results in greater values of explained variance statistics: these are R-squared values in the case of our simulations. This is because statistical models suffering from this bias explain both true and artificial (due to collider) variation in a dependent variable as we show in the Supplementary Data. This further complicates the assessment of the relative predictive power of polygenic scores and environments. As demonstrated in Figure 1C, R-squared statistics for the model with an endogenous covariate and a polygenic score would include not only the true share of the variance in Y explained by G and E, but also the elements of variance that are due to rGE and confounder(s), U.

To conclude, the inclusion of associated polygenic scores and covariates in regression models may result in spurious estimates and greater explained variance statistics. The direction and strength of coefficient bias depends on the strength of the gene-covariate correlation and on the underlying structure of any endogeneity which links the covariate to the outcome variable.

### Gene × environment interaction models

A growing literature estimates the moderating patterns of environmental risks in the associations between polygenic scores and phenotypes. Here, in the same fashion as in additive models, environmental exposures of interest are not usually exogenous. For example, recent studies on gene-environment interaction analysis consider such environments as relationship status,^22^ educational attainment,^23^ occupational exposure,^24^ neighbourhood characteristics^25^ and others. There is ongoing discussion on the implications of non-exogeneity of environments.^7; 26; 27^ Also, the issue of collider bias has been demonstrated in the context of case-only gene-environment interaction studies.^28^ We expand on these concerns by showing that moderation models also suffer from collider bias.

Firstly, the problem outlined until this point is also relevant for gene-environment interaction studies. One difference, however, between additive and moderation models is the presence of the G×E interaction in the set of covariates. As indicated in Figure 2A by green nodes, the bias path from G to Y through E and U would still lead to spurious results. Considering the examples of G×E studies mentioned earlier, the environments may, to some degree, be products of self-selection, which leads to a greater likelihood of G and E interdependence along with the presence of unobserved confounder(s), U. Secondly, since the overall G×E interaction pattern depends on the direct estimates of G on Y and E on Y, results for moderation analyses are biased when direct effects are spurious. However, the G×E coefficient *per se* is not inflated due to collider bias.^29^ This can be seen in Figure 2B, along with the inflation of R-square statistics in Figure 2C which were obtained from similar simulations as earlier but with the inclusion of interaction terms. Consequently, endogeneity of environmental covariates biases both additive and moderation models.

**Figure 2.**
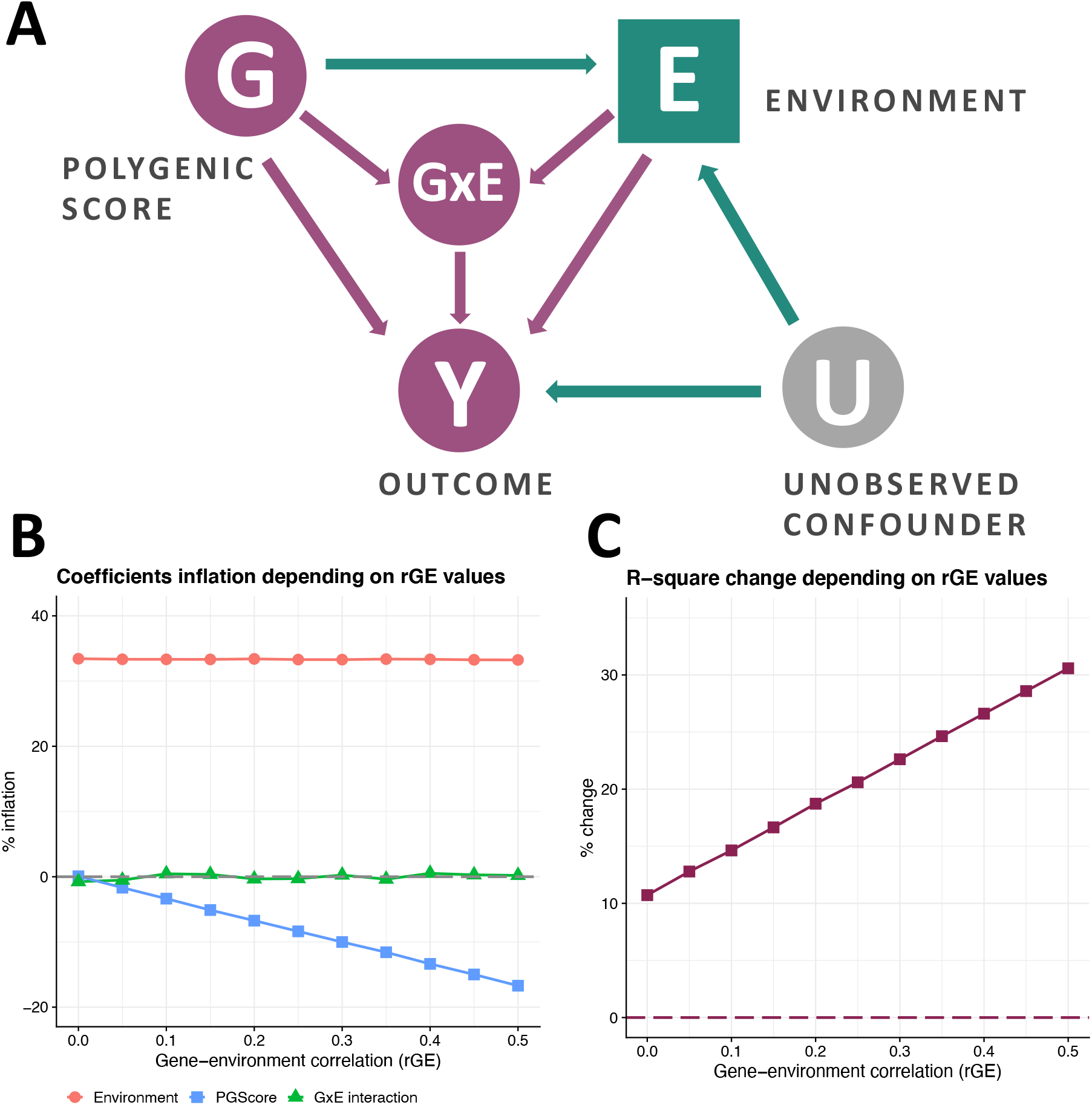
Collider bias in polygenic gene-environment interaction models. Panel A. Schematic diagram of the collider bias which occurs between polygenic score, environment, and outcome in cases of gene-environment interdependence. Dark purple circles represent variables, unobserved confounders are shown in grey circles, collider variables are indicated in squares. By adding E into the model with the polygenic score G, we make E a collider. A collider that is not conditioned on, blocks the path between its sources (G and U); once a collider is controlled for, the path is opened as indicated by green nodes. Panel B. Spurious regression estimates for the polygenic score and environment along with non-inflated interaction terms from the series of OLS simulations reflecting a range of gene-environment interdependence. Collider bias due to positive values of gene-environment correlation and the presence of an uncontrolled confounder, which is positively correlated with covariate and outcome, results in deflation of polygenic score estimates. Deflation is greater the higher the gene-environment correlation; greater confounding also results in greater bias. The interaction term is not affected but results for moderation analysis are biased as long as direct effects are spurious. Panel C. R-squared inflation plot from the series of OLS simulations; collider bias results in inflated values of explained variance statistics. R-squared statistics for the model with endogenous covariate and polygenic score includes not only the true share of the variance in Y explained by G and E (baseline estimate indicated by 0), but also the elements of variance that are due to gene-environment correlation and confounder(s), U.

### Solutions

To address this issue, it is important to understand the nature of the correlation between genes and environment (or phenotype if it is used as a covariate) – whether a correlation is conditional and observed because of omitted confounders between G and E and/or a correlation reflects causal dependency. The former would necessitate controlling for the confounding factor: this could be parental characteristics (passive rGE)^30; 31^, ancestry^2^, and so forth. If a correlation arises as a result of active or evocative selection, the assumption of non-causal association would be violated and require another set of solutions to avoid collider bias. The latter is relevant when a phenotypic variable is used as a covariate along with its polygenic prediction, since the association would be at least partially causal.

To obtain unbiased estimates, we need to apply causal inference methods that seek to exploit the exogenous variation in an environmental covariate. A comprehensive discussion of methods available for researchers and applicable to the context of this paper is provided by Fletcher and Conley.^6^ Briefly, techniques such as regression discontinuity and difference-in-difference designs, instrumental variables and quasi-natural experiments will produce unbiased results for both additive and gene-environment interaction models, conditional on certain assumptions being met.

There are also existing ways to assess the magnitude of the bias for the general collider cases.^32; 33^ Since the type of collider we described here is the product of both observed and unobserved factors, calculation of bias magnitude would rely on additional assumptions about the structure of the error correlation between the environmental covariate and the outcome. The presence of unobserved variables, however, makes it impossible to provide a definitive estimate of the bias. However, in the Supplementary Data we show the sources of bias in coefficient estimates and error in R-squared. We demonstrate that the strength of collider bias is positively associated with the strength of rGE and unobserved confounders. The use of sensitivity analyses is a valuable tool in showing how robust conclusions are to different degrees of unobserved confounding and thus of collider bias.^34^

## Conclusion

We have discussed methodological considerations arising due to heritable environments (or phenotypes that are included into models as covariates) that have not yet been previously recognised. We demonstrated that the inclusion of environments that are products of polygenic scores may introduce endogenous selection bias through conditioning on a collider, leading to spurious associations. Particularly, we showed that the degree of bias depends on the strength of the gene-covariate correlation and of the omitted variable(s) linking the covariate and outcome. We also showed that the portion of explained variance is overestimated proportionally. We proposed some solutions that exploit the strength of causal inference methods: these are likely to be important not only for obtaining reliable results but also in the interpretation of existing studies.

## Supporting information

Bias in gene-environment models due to interdependency of polygenic scores and environments

## Supplemental Data

Supplemental Data include the detailed derivations for the bias under the assumption of linear relationships that are modelled using regression analysis.

## Acknowledgments

Research has been supported by the UKRI/ESRC NCRM SOCGEN grant, ERC Advanced Grant (CHRONO, 835079) and The Leverhulme Trust Large Centre Grant, Leverhulme Centre for Demographic Research (PI, M.C. Mills). Results from this research were presented earlier at the National Institute on Aging supported 2019 Integrating Genetics and Social Science Conference (R13-AG062366) at the University of Colorado, Boulder.

## Declarations of Interest

The authors declare no competing interests.

## Data and Code Availability

The code for the simulations and figures is available on Zenodo (DOI: 10.5281/zenodo.4184673) and GitHub (https://github.com/eva-akimova/collider-simulations.git).

## Notes

### Competing Interest Statement

The authors have declared no competing interest.

