## Supplementary material for "Heritable environments: bias due to conditioning on a collider in models with polygenic scores": Bias in gene-environment models due to interdependency of polygenic scores and environments

### Supplemental Data: Bias in gene-environment models due to interdependency of polygenic scores and environments

The main text provided a general description of endogenous selection bias when polygenic scores and environments are not independent. Here we further illustrate this issue by deriving exact expressions for the bias under the assumption of linear relationships that can be modelled using regression analysis.

We assume that the data have been generated by the DAG shown in Figure 1A. Here U is an unobserved variable or set of variables that confounds the E – Y relationship (this is equivalent to, but, in our view more transparent than, a depiction that would include correlated error terms for E and Y). We further assume that all the variables have unit standard deviation and that G is exogenous.

*S1.1 Additive model*

When the effects of G and E on Y are additive the true linear models given the data-generating process are:

$$E(E|G,U) = \alpha_0 + \alpha_1 G + \alpha_2 U \qquad \text{(Equation S1)}$$

$$E(Y|G,E,U) = \beta_0 + \beta_1 G + \beta_2 E + \beta_3 U \qquad \text{(Equation S2)}$$

We assume that the parameters $\alpha_1$, $\beta_1$, and $\beta_2$ are all positive.

If estimate the models:

$$E(E|G) = a_0 + a_1 G \qquad \text{(Equation S3)}$$

$$E(Y|G,E) = b_0 + b_1 G + b_2 E \qquad \text{(Equation S4)}$$

the relationships between the true and estimated parameters are:

$$b_1 = \beta_1 - \frac{\alpha_1 \alpha_2 \beta_3}{1-\alpha_1^2} \qquad \text{(Equation S5)}$$

$$b_2 = \beta_2 + \frac{\alpha_2 \beta_3}{1-\alpha_1^2} \qquad \text{(Equation S6)}$$

The proof is as follows. Let $\beta_{YG}$ denote the coefficient of the unconditional regression of Y on G and likewise for $\beta_{YE}$. Then, tracing the paths linking G and Y in the Figure A1a we have:

$$\beta_{YG} = \beta_1 + \beta_2 \alpha_1$$

and

$$\beta_{YE} = \beta_2 + \beta_1 \alpha_1 + \beta_3 \alpha_2$$

Given that $\beta_{EG} = \alpha_1$ we then apply the standard formula to derive conditional regression coefficients from unconditional:

$$b_1 = \frac{\beta_{YG} - \beta_{YE}\alpha_1}{1-\alpha_1^2} = \frac{\beta_1 + \beta_2\alpha_1 - (\beta_2 + \beta_1\alpha_1 + \beta_3\alpha_2)\alpha_1}{1-\alpha_1^2}$$

Straightforward algebra yields (S5). $b_2$ is derived similarly. Notice, however, that $a_1 = \alpha_1$ because G and U are unconditionally independent.

The bias in both estimates depends on the sign of $\alpha_2 \times \beta_3$: if this is positive the estimate of the partial effect of G on Y, given E, will be downwardly biased and the estimate of the effect of E on Y, given G, will be upwardly biased. If there is no correlation between G and E ($\alpha_1 = 0$) then $b_1$will be unbiased. If there is no effect of an unmeasured confounder (either $\alpha_2 = 0$ and/or $\beta_3 = 0$) both $b_1$ and $b_2$ will be unbiased. The bias in the effect of G on Y has a different sign than the bias in the effect of E on Y: if the bias in the latter is positive, the size of the genetic effect will be underestimated relative to the environmental effect.

The example of coefficient deflation from Papageorge and Thom[1] (Table 6: 41) can be demonstrated following Equation S5. For instance, considering the case on nonroutine interactive job tasks as the dependent variable, we see that the baseline coefficient of the educational attainment polygenic score is 0.185, which reflects a model without any environmental and phenotypic covariates. In the model with educational controls (respondent's years of schooling and parental education), the polygenic score coefficient drops to 0.055 reflecting a 70% negative change. Since the dependent variable is standardised, we can assess the relative importance of collider bias which is $\frac{\alpha_1\alpha_2\beta_3}{1-\alpha_1^2}$ from Equation S5 under additional assumptions. If we allow the coefficient of the correlation between educational attainment polygenic score and respondents years of schooling $\alpha_1$=0.300, and the presence of unobserved confounder U, positively correlated with both years of schooling and job task (for example, living in advantaged neighbourhood as a child), we have $\alpha_2 = 0.250$ and $\beta_3 = 0.250$. These are all plausible and rather modest suggestions following correlation matrix from Table 6, leading the inflation bias to be:

$$\frac{\alpha_1\alpha_2\beta_3}{1-\alpha_1^2} = \frac{0.300 \times 0.250 \times 0.250}{1-0.300^2} = 0.021$$

which explains 16% downward change of polygenic score coefficient.

*S1.2 G×E interaction model*

The DAG in Figure 2A shows the case in which the effect of G on Y varies with E. In this case, the true linear models given the data-generating process are:

$$E(E|G,U) = \alpha_0 + \alpha_1 G + \alpha_2 U \quad \text{(Equation S7)}$$

$$E(Y|G,E,U) = \beta_0 + \beta_1 G + \beta_2 E + \beta_3 U + \beta_4 (GE) \quad \text{(Equation S8)}$$

We estimate:

$$E(E|G) = a_0 + a_1 G \quad \text{(Equation S9)}$$

$$E(Y|G,E) = b_0 + b_1 G + b_2 E + b_4 GE \quad \text{(Equation S10)}$$

In this case, $b_4$ is an unbiased estimate of $\beta_4$ because the backdoor path from G-E to Y is blocked by E. The bias in $b_1$ and $b_2$ will be the same as above. In the case in which E is a binary variable, coded 0 and 1, $b_4$ will be an unbiased estimate of the difference in the effect of G at $E = 1$ and $E = 0$, but the estimate of the baseline effect of G on Y when $E = 0$ will be biased.

*S1.3 Bias in $R^2$*

The $R^2$ for models S4 and S10 will be biased. In the additive case, for example, the true $R^2$ attributable to G and E is:

$$\frac{\beta_1^2 var(G) + \beta_2^2 var(E) + 2\beta_1\beta_2 cov(G,E)}{var(Y)} = \beta_1^2 + \beta_2^2 + 2\beta_1\beta_2\alpha_1 \quad \text{(Equation S11)}$$

(using the assumption that all the variables have unit standard deviation). But the reported $R^2$ from model S4 is:

$$b_1^2 + b_2^2 + 2b_1b_2\alpha_1 \qquad \text{(Equation S12)}$$

Substituting S5 and S6 into S12 we calculate the inflation of $R^2$ due to confounding and collider bias. This is:

$$R^2bias = \alpha_2\beta_3\left[\frac{\alpha_2\beta_3}{1-\alpha_1^2} + 2\beta_2\right] \qquad \text{(Equation S13)}$$

Confounder bias arises from $\alpha_2\beta_3$. The derivative of S13 with respect to this is positive provided that $1 - \alpha_1^2 > 0$. The derivative of S13 with respect to $\alpha_1$ (which captures the association between G and E) is:

$$\frac{2\alpha_1\alpha_2^2\beta_3^2}{(1-\alpha_1^2)^2}$$

The sign of this depends on the sign of the numerator. When it is positive both confounding and collider bias will inflate the reported $R^2$. As an example, consider a case in which $\beta_1 = 0.465, \beta_2 = 0.505, \beta_3 = 0.231, \alpha_1 = 0.209, \alpha_2 = 0.693$. Then the observed $R^2$ = 0.758, whereas the true share of the variance in Y explained by G and E is 0.569. The inflation bias here is:

$$0.693 \times 0.231\left[\frac{0.693 \times 0.231}{1 - 0.209^2} + 2 \times 0.505\right] = 0.188$$

If the correlation between G and E had been larger and/or if the confounding of E had been greater, the reported $R^2$ would have been larger because of the greater bias.
